# Effects of Heat Shock and Smoke on the Germination of Woody Species from Tropical Dry Forests

**DOI:** 10.64898/2026.10.04.756604

**Authors:** Natalia Andrea Díaz Bernal, María Clara Castellanos, Ángela Parrado Rosselli

**Author notes:** **Corresponding author:** Natalia Andrea Díaz Bernal.

## Abstract

Fire-dependent or fire-influenced ecosystems harbor species with diverse response strategies, such as smoke-stimulated germination and dormancy release through heat shocks. Although these cues influence germination in many fire-prone ecosystems, their effects on seeds of woody species from tropical dry forests (TDFs) remain poorly understood. We evaluated the germination response of twelve TDF woody species to three heat-shock temperatures (80, 120, and 160°C), two exposure times (3 and 5 min), and two smoke conditions (with and without smoke water), plus dry and wet controls. We also assessed the relationship between germination and functional seed traits, including permeability, seed-coat thickness, mass, and shape index. Of the eleven species that germinated, four showed broad tolerance, with no significant reduction in germination even at 160°C for 5 min, which was associated with lower permeability, thicker seed coats, and greater mass. Four other species showed restricted tolerance to heat, with significant declines in germination at higher temperatures or longer exposure times, and were associated with higher permeability, thinner seed coats, and lower seed mass. The remaining three species showed heat-stimulated germination between 80 and 120°C and tended to have lower shape-index values. Smoke treatments significantly increased germination in three species, *Ceiba pentandra, Guazuma ulmifolia*, and *Samanea saman*. In the remaining species, significant differences were either absent or attributable to moisture or imbibition effects rather than to smoke itself. Overall, germination responses to heat were associated with variation in functional seed traits, whereas responses to smoke were less widespread and showed a less consistent functional pattern.

## Introduction

Tropical dry forests (TDFs) are among the most threatened ecosystems and a top conservation priority worldwide (Miles et al. 2006; Karna and Jayakumar 2026). In Colombia, TDFs are critically endangered, mainly due to landscape transformation into cattle pastures, agricultural fields, and urban areas (García et al. 2014; Pizano et al. 2014; Etter et al. 2017; González-M et al. 2018; Díaz-Triana et al. 2019). As a result, less than 8% (720,000 ha) of their original cover remains, distributed mainly in vegetation mosaics at different successional stages, embedded within a landscape matrix dominated by livestock and agricultural activities (García et al. 2014). In these landscapes, the combination of marked dry seasons, fuel accumulation from deciduous species, and the introduction of flammable invasive grasses and shrubs (Valderrama et al. 2018; Karna and Jayakumar 2026), together with the recurrent use of fire in agricultural practices (Armenteras et al. 2020; Ramírez 2020), contributes to high fire activity in these ecosystems (Stan and Sánchez-Azofeifa 2019; Karna and Jayakumar 2026). However, the responses of TDF plant species to fire remain poorly understood.

In ecosystems where fire is a natural disturbance, it can favor key ecological processes such as resprouting, serotiny, and germination stimulated by fire-related cues, including heat shock and smoke (Bradshaw et al. 2011; Keeley et al. 2011; Pausas and Keeley 2014; Keeley and Pausas 2022). This last process occurs when seeds exposed to high temperatures or combustion by-products show greater germination success than in the absence of these stimuli (Keeley and Fotheringham 2000; Paula and Pausas 2008; Pausas and Keeley 2014; Pausas and Lamont 2022). Heat shock can modify the seed coat through scarification, desiccation, or weakening of the testa, thereby facilitating imbibition and promoting germination (Keeley and Fotheringham 2000; Moreira and Pausas 2012; Keeley and Pausas 2022; Pausas and Lamont 2022). Smoke, in turn, acts as a chemical cue through compounds such as karrikinolides, promoting physiological processes associated with dormancy release (Van Staden et al. 2000; Light et al. 2009; Soós et al. 2019; Pausas and Lamont 2022).

Although heat and smoke are recognized as the main drivers of fire-related germination (Keeley and Fotheringham 2000; Pausas and Lamont 2022), germination responses depend not only on the individual action of these cues but also on how they interact with each other (Thomas et al. 2003, 2007; Soós et al. 2019; Zirondi et al. 2019; Cuello et al. 2020; Hodges et al. 2021). Under natural conditions, seeds may be exposed to heat during a fire and subsequently to chemical compounds present in smoke (Pausas 2012; Pausas and Lamont 2022). Therefore, evaluating both cues independently may not fully represent the conditions to which seeds are exposed after a fire (Thomas et al. 2003; Pausas 2012). In addition, the response to their joint action varies among species: in some it may reduce germination, whereas in others it may generate additive or synergistic effects relative to the individual action of heat or smoke (Thomas et al. 2003, 2007; Moreira et al. 2010; Zirondi et al. 2019; Hodges et al. 2021; Pausas and Lamont 2022).

This variation in seed responses to heat and smoke is largely related to variation in functional seed traits such as permeability, seed-coat thickness, mass, and shape index, which may influence the ability of species to tolerate or be stimulated by disturbance or, conversely, to be sensitive to its effects (Gómez-González et al. 2011; Enright et al. 2014; Ooi et al. 2014; Daibes et al. 2019; Hradilová et al. 2019; Zirondi et al. 2019; Newton et al. 2021; Lamont et al. 2022; Pausas and Lamont 2022). For example, seed-coat permeability and thickness influence embryo protection and regulate water uptake and the onset of germination (Mohamed-Yasseen et al. 1994; Wiggers et al. 2017; Lamont et al. 2022). Impermeable seeds may require high temperatures to break dormancy, whereas smoke may stimulate germination in permeable seeds (Paula and Pausas 2008; Moreira et al. 2010; Zirondi et al. 2019; Pausas and Lamont 2022). Similarly, greater seed mass and a more compact (rounder) shape, which minimizes the surface-area-to-volume ratio, may slow heat transfer to the embryo, whereas lighter seeds that are more elongated or flattened may be more sensitive to high temperatures (Tavşanoğlu et al. 2017; Daibes et al. 2019; Moro et al. 2023).

In Colombia, the few studies conducted to date have shown contrasting responses among woody species: some tolerate heat or show increased germination at certain temperatures, whereas others lose viability under more intense heat shocks (Cárdenas-Salgado and Pizano 2019; Hernández 2021; Barras et al. 2025). However, these studies have focused primarily on heat shock, leaving the role of smoke, its interaction with heat, and the extent to which functional traits explain interspecific variation in the germination responses of woody species from Colombian tropical dry forests largely unexplored. Understanding seed tolerance and the functional traits associated with it could inform post-fire restoration in tropical dry forest landscapes subject to recurrent fires. Such information could help identify species capable of persisting under disturbance in human-modified landscapes, as well as those that may require enrichment actions to maintain ecosystem functional diversity.

In this context, we evaluated the effects of heat shock and smoke on the germination of TDF woody species and their relationships with seed permeability, seed-coat thickness, mass, and shape index. We first hypothesized that the effects of combined heat and smoke treatments on seed germination would exceed those of heat shock alone (Thomas et al. 2003, 2007; Monemizadeh et al. 2025). We further hypothesized that seeds with low permeability, thick seed coats, greater mass, and a more compact shape (lower shape index) would show greater tolerance to heat shock and retain their germination capacity after exposure to high temperatures. In contrast, we expected more permeable seeds with thinner seed coats, lower mass, and more elongated or flattened shapes (higher shape index) to respond positively to smoke treatments and low-intensity heat shocks (Moreira et al. 2010; Ribeiro et al. 2015; Fichino et al. 2016; Daibes et al. 2019; Hradilová et al. 2019; Monemizadeh et al. 2025).

## Materials and methods

### Study area and seed collection

We sampled four representative tropical dry forest (TDF) sites in the upper Magdalena River basin, in Honda, Tolima, Colombia: La Picota (5°13’26.22" N, 74°47’41.30" W), El Tambor (5°11’00.35" N, 74°47’22.37" W), El Triunfo (5°10’06.75" N, 74°45’48.21" W), and San Jorge (5°13’04.1" N, 74°46’02.7" W). Species were selected based on their representativeness of Colombian tropical dry forests, following Pizano and García (2014) and Aguilar et al. (2016). We also sought variation in functional traits, although the final selection was constrained by the availability of viable fruits and seeds during the collection period (Table 1).

**Table 1.** Woody species selected for the germination experiments

| Family | Species | Dormancy type | Successional stage |
| --- | --- | --- | --- |
| Fabaceae | *Machaerium capote* | Non-dormant (Morales-Pisco and Parrado-Rosselli unpubl.) | Intermediate to late secondary (Avella-M et al. 2019) |
| Zygophyllaceae | *Bulnesia carrapo* | Non-dormant (Pañitrur et al. 2021) | Intermediate to late secondary (Pañitrur et al. 2021) |
| Fabaceae | *Pithecellobium dulce* | Physical (Singh et al. 2021) | Early secondary (Avella-M et al. 2019) |
| Malvaceae | *Ceiba pentandra* | Physical (Egbewole and Clement 2016) | Pioneer / early secondary (Ribeiro et al. 2023) |
| Anacardiaceae | *Anacardium excelsum* | Physical (Salcedo et al. 2017) | Intermediate secondary (Yepes-Quintero et al. 2007) |
| Fabaceae | *Chloroleucon mangense* | Physical (Rodríguez-Trejo and Guizar-Nolazco 2016) | Pioneer / early secondary (Avella-M et al. 2019) |
| Fabaceae | *Enterolobium cyclocarpum* | Physical (Peraza-Villarreal et al. 2018) | Early to intermediate secondary (Yepes and Villa 2010) |
| Fabaceae | *Pseudosamanea guachapele* | Physical (Labre et al. 2022) | Pioneer / early secondary (Castañeda-Garzón et al. 2024) |
| Salicaceae | *Casearia corymbosa* | Non-dormant (Morales-Pisco and Parrado-Rosselli unpubl.) | Early to intermediate secondary (Avella-M et al. 2019) |
| Malvaceae | *Guazuma ulmifolia* | Physical (Sánchez et al. 2017) | Pioneer / early secondary (Avella-M et al. 2019) |
| Fabaceae | *Samanea saman* | Physical (Peraza-Villarreal et al. 2018) | Early to intermediate secondary (Cascante et al. 2002) |
| Rutaceae | *Zanthoxylum rhoifolium* | Physical and physiological (Corrêa et al. 2022) | Intermediate secondary (Avella-M et al. 2019) |

Fruits and seeds were collected during 2025 following the guidelines in the wild plant seed collection manual by Di Sacco et al. (2020). Sampling took place during the transition into the first and second rainy seasons, except for *Guazuma ulmifolia*, which was sampled during the mid-year dry season (Online Resource 1). Fruits were collected when species showed signs of maturity or natural dispersal, based on morphological indicators such as color change, fruit drying, natural opening of dehiscent fruits, or the presence of recently dispersed seeds. For each species, material was collected from at least ten randomly selected individuals separated by a minimum distance of 30 m.

Collection was carried out primarily from parent trees by manually harvesting accessible fruits or cutting fruit-bearing branches. For *Ceiba pentandra*, *Anacardium excelsum*, *Pseudosamanea guachapele*, and *Samanea saman*, recently fallen fruits or seeds were also collected from the ground. In these cases, only intact, recently dispersed material with no visible signs of deterioration, predation, or fungal attack was selected.

In the field, fruits were placed in paper bags, and seeds were extracted on the same day as collection to prevent deterioration. The seeds were then transported to the laboratory and dried under ventilated conditions. Once dry, they were stored in airtight glass jars in a cool place protected from direct light for a maximum of six months (Online Resource 1).

### Functional seed traits

Functional seed traits were characterized using seeds from the same batches used in the germination experiments. Four traits were measured across the twelve study species: permeability, seed coat thickness, mass, and shape index, using 50 randomly selected seeds per species (Online Resource 2).

Seed coat permeability was assessed through imbibition tests, in which seeds were submerged in distilled water for 24, 48, and 72 h. After each imbibition interval, the percentage increase in mass relative to initial mass was calculated. Seeds were classified as water permeable when the maximum increase observed at any of the three time points exceeded 20% (Moreira et al. 2010; Zirondi et al. 2019). Seed coat thickness was measured from transverse seed sections, and the resulting images were analyzed using ImageJ software.

Seed mass was determined by weighing each seed individually on an analytical balance with a resolution of ± 0.0001 g, after oven drying at 60 °C for 72 h (Pérez Harguindeguy et al. 2013; Daibes et al. 2019). Finally, shape index was calculated following Pérez Harguindeguy et al. (2013). Three dimensions were measured for each seed: length, width, and thickness, and expressed in millimeters. Each dimension was divided by the largest of the three measurements recorded for that seed, after which the variance of the three normalized values was calculated. The shape index ranged from 0 to 1, with values close to 0 indicating more spherical seeds and values close to 1 indicating more elongated or flattened seeds (Thompson et al. 1993; Pérez Harguindeguy et al. 2013; Moro et al. 2023). Because the index does not distinguish between elongated and flattened seeds, we refer to high values as “elongated or flattened.”

### Heat shock treatments

Seeds were subjected to six heat shock treatments resulting from the combination of three temperatures (80, 120, and 160 °C) and two exposure times (3 and 5 min), together with a dry control that was not exposed to heat. The number of replicates and the number of seeds per replicate were defined with reference to previous studies evaluating germination responses to heat shock (for example, Daibes et al. 2019; Zomer et al. 2022; Barras et al. 2025). Accordingly, five independent replicates were established for each treatment, and the number of seeds per replicate ranged from 6 to 50 depending on the seed size and availability of each species (Online Resource 1). Treatments were conducted in a preheated oven under controlled temperature conditions, and each replicate was treated separately to ensure sample independence.

Treatment selection was based on previous experimental studies conducted in Colombian tropical dry forests and other tropical dry ecosystems, where germination and viability responses have been evaluated under heat shocks ranging from 60 to 200 °C and exposure times of up to 10 min (Jaureguiberry and Díaz 2015; Ribeiro et al. 2015; Fichino et al. 2016; Oliveira et al. 2018; Cárdenas- Salgado and Pizano 2019; Daibes et al. 2019; Zirondi et al. 2019; Hernández 2021; Moro et al. 2021; Barras et al. 2025). These treatments also approximate the thermal conditions that seeds may experience during surface fires in tropical dry forests. Such fires primarily consume leaf litter and dead organic material available on the forest floor and are generally classified as events of low to moderate intensity (Karna and Jayakumar 2026). In this context, 80 °C represented moderate thermal exposure consistent with temperatures recorded near the soil surface under high fuel loads in the Brazilian Cerrado, whereas 120 and 160 °C allowed us to evaluate responses to more severe or extreme thermal conditions (Daibes et al. 2017; Zupo et al. 2020).

### Heat shock and smoke treatments

Exposure to smoke-derived compounds was achieved by imbibing the seeds in smoke water, hereafter referred to as the smoke treatment. Following Jäger et al. (1996) and Moreira et al. (2010), a solution was prepared from plant biomass composed of leaves and fine branches collected from the study area in the tropical dry forest of Honda, Tolima. The biomass was dried, ground, and subsequently exposed to 200 °C for 30 min. Distilled water was then added at a ratio of 10 mL per gram of dry biomass, and the mixture was allowed to stand for at least 30 min. Finally, the solution was filtered and used at a 1:1 concentration.

To evaluate the combined effect of heat shock and smoke, six treatments were established from the combination of three temperatures, 80, 120, and 160 °C, and two exposure times, 3 and 5 min. In each case, seeds were first exposed to the heat shock treatments and subsequently imbibed in smoke water for 24 h. Each combination was evaluated using five replicates. The wet control consisted of seeds imbibed only in distilled water for the same period.

### Germination assay

After treatment exposure, seeds were placed in Petri dishes lined with two layers of filter paper moistened with distilled water and incubated in a germination chamber at 28 °C, corresponding to the mean annual temperature reported for Colombian tropical dry forests (González-M et al. 2018), under a 12 h light and 12 h dark photoperiod. Germination was monitored for two months. During the first month, assessments were conducted every two days, and during the second month, every eight days. Throughout this period, seeds were moistened as needed to maintain suitable germination conditions. Seeds were considered germinated when radicle emergence was observed.

### Viability tests

Before treatment application, and depending on seed availability, viability tests were conducted following an adaptation of the ISTA (2025) guidelines. Three replicates of 20 seeds per species were used. Seeds were imbibed in distilled water at 20 °C for 24 h. For species with impermeable seed coats, seeds were scarified beforehand with 250 grit sandpaper to facilitate imbibition. After this pretreatment, a longitudinal cut was made through the embryo, and the seeds were incubated in a 1% solution of 2,3,5 triphenyltetrazolium chloride (TTC) at 30 °C. To determine the most appropriate staining time for each species, intervals of 0.5, 1, 2, 3, and 4 h were evaluated based on individual observations of embryo staining patterns.

At the end of the two-month germination monitoring period, ungerminated seeds that had not lost their firmness were assessed using the viability test under the conditions previously established for each species. Final seed viability was calculated as the number of viable ungerminated seeds plus the number of seeds that germinated during the experiments.

### Statistical analysis

All statistical analyses were conducted in R version 4.5.2 (R Core Team 2025). First, differences in germination probability between the dry control and the heat shock treatments were evaluated using generalized linear models (GLMs) for each species. For each species, we also explored the inclusion of replicate as a random effect using generalized linear mixed models (GLMMs). Because the estimated variance associated with this effect was close to zero, the mixed models did not provide additional information beyond the GLMs. Therefore, the final analyses were conducted using GLMs, with treatment included as a fixed effect.

For these models, a binomial distribution and Firth-penalized regression were used, implemented through the brglm2 package (Kosmidis 2025). This approach was used to reduce problems associated with complete or quasi-complete separation in the data, following the finiteness and bias-reduction properties described for penalized binomial models (Kosmidis and Firth 2021). For *Ceiba pentandra*, a quasibinomial GLM with a Haldane-Anscombe correction was fitted because of the observed overdispersion. Estimated marginal means and contrasts between the dry control and the treatments were obtained using the emmeans package (Lenth and Piaskowski 2025). Comparisons of each heat-shock treatment and the wet control with the dry control were performed using estimated marginal means contrasts, with *p*-values adjusted for multiple comparisons using Dunnett’s method within each species (Dunnett 1955).

To evaluate the effect of smoke, we first assessed whether germination responses could be influenced by the moisture associated with smoke water. For this purpose, germination was compared between the dry control and the wet control. When this comparison was significant, the difference was attributed to the effect of moisture or imbibition rather than to smoke. A second set of generalized linear models was then fitted to compare germination responses between heat shock treatments alone and combined heat shock and smoke treatments under the same temperature and exposure time. As in the previous analysis, Firth-penalized regression was used through the brglm2 package. For *Ceiba pentandra*, a quasibinomial model with a Haldane-Anscombe correction was fitted. Pairwise comparisons between treatments with and without smoke were performed using emmeans.

Validation of the binomial models fitted with Firth-penalized regression was conducted through residual simulation using the DHARMa package (Hartig 2024). Assumptions related to uniformity, dispersion, outliers, and residual patterns were examined. For *Ceiba pentandra*, validation of the quasibinomial model was based on inspection of overdispersion, residual behavior, and the presence of influential observations.

Finally, to explore the relationship between germination responses and functional seed traits, a principal component analysis (PCA) was conducted. Four standardized traits were included in this analysis: permeability, seed coat thickness, mass, and shape index. PC1 and PC2 scores were initially evaluated in relation to germination response. Because PC2 showed no significant relationships with log(OR) across treatments, subsequent analyses and interpretation focused on PC1.

Based on the previously fitted generalized linear models, the odds ratio (OR) of germination was estimated for each species and heat treatment relative to its corresponding dry control. For the effect of smoke, ORs were calculated from pairwise comparisons between each heat treatment with smoke and its equivalent treatment without smoke under the same temperature and exposure time. The resulting OR values were transformed using the natural logarithm. Independent linear models were then fitted for each temperature and exposure time combination, using log(OR) as the response variable and PC1 as the explanatory variable. Model validation included assessment of residual normality using the Shapiro-Wilk test and identification of potentially influential observations using Cook’s distance.

## Results

### Effects of heat shock on seed germination

Of the 12 species evaluated, 11 germinated in at least one treatment, whereas *Zanthoxylum rhoifolium* did not germinate under any of the conditions evaluated. Among the species that germinated, tolerant and heat-stimulated germination responses were identified (Fig. 1). Four species showed restricted tolerance, four showed broad tolerance, and three showed heat-stimulated germination. Restricted tolerance included species whose germination was similar to the control only under moderate heat shocks but decreased significantly as temperature or exposure time increased. Broad germination tolerance included species that showed no significant reductions in germination relative to the control in any treatment, even at 160 °C for 5 min. Finally, heat-stimulated germination corresponded to species showing significant increases relative to the control in at least one heat shock treatment.

**Fig. 1.**
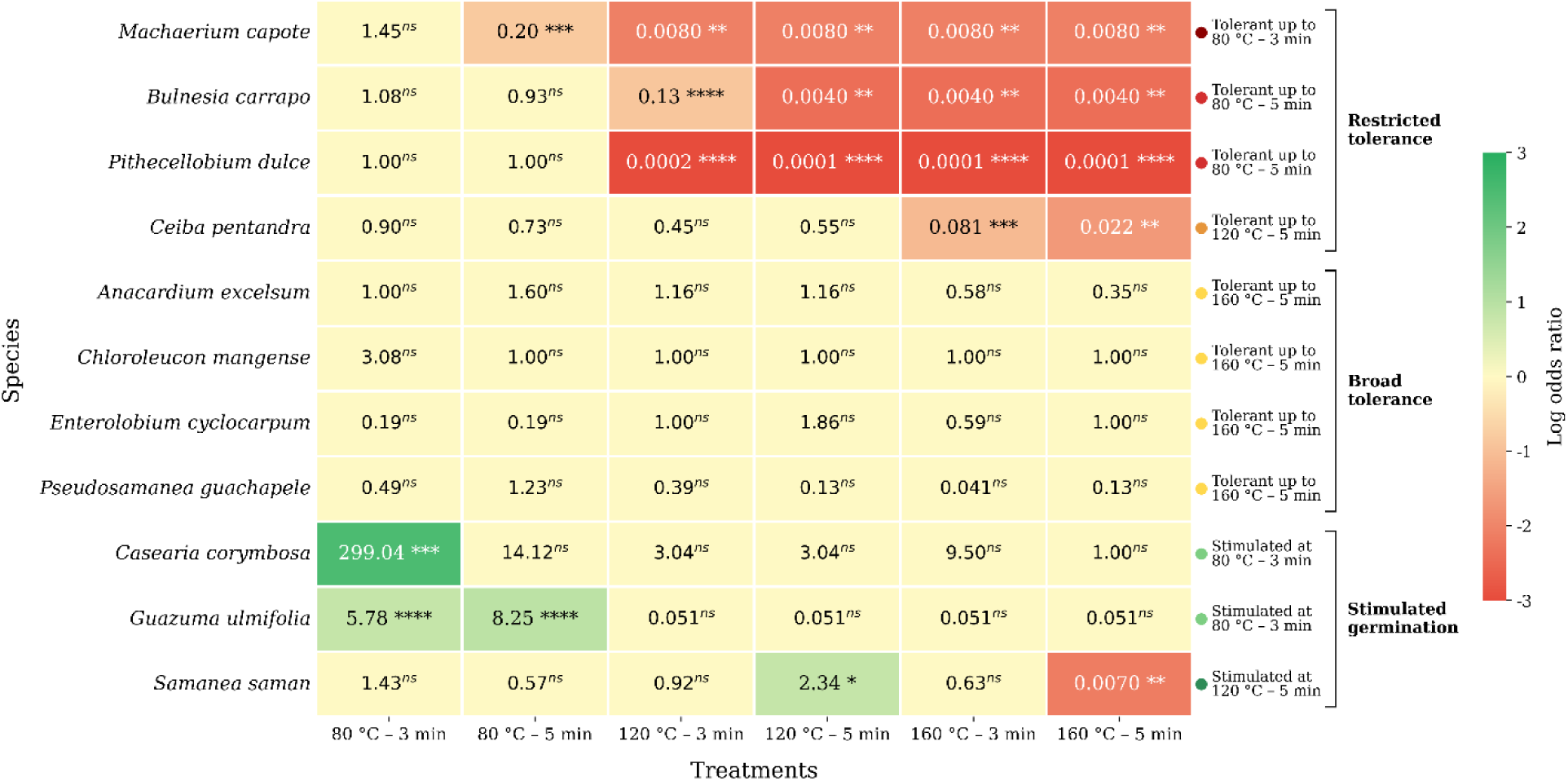
Heatmap of odds ratios for the probability of germination of 11 species evaluated under heat shock treatments relative to the dry control. Each cell shows the estimated odds ratio for each treatment, represented on a log10 color scale: green tones indicate a higher probability of germination, whereas red tones indicate a lower probability. Treatment codes indicate temperature and exposure time (e.g., T80–3 = 80 °C for 3 min). ns = not significant; * P ≤ 0.05; ** P ≤ 0.01; *** P ≤ 0.001; **** P ≤ 0.0001

Among the species with restricted tolerance, *Machaerium capote* tolerated heat shock up to 80 °C for 3 min. Germination decreased from 37% in the control to 10% at 80 °C for 5 min, with a significant reduction in germination probability (OR = 0.20, *p* ≤ 0.001) (Table 2; Online Resource 3). *Bulnesia carrapo* and *Pithecellobium dulce* showed tolerance up to 80 °C for 5 min. In both species, germination decreased significantly from 120 °C for 3 min onward compared with the control. In *B. carrapo*, germination decreased from 66.7% in the control to 20.0%, whereas in *P. dulce* it decreased from 98.4% to 0.8%. In both cases, germination probability was significantly lower (OR = 0.13, *p* < 0.0001; OR = 0.0002, *p* < 0.0001, respectively). *Ceiba pentandra* tolerated heat shocks up to 120 °C for 5 min (*p* > 0.05), after which germination decreased significantly from 160 °C for 3 min onward.

**Table 2.**
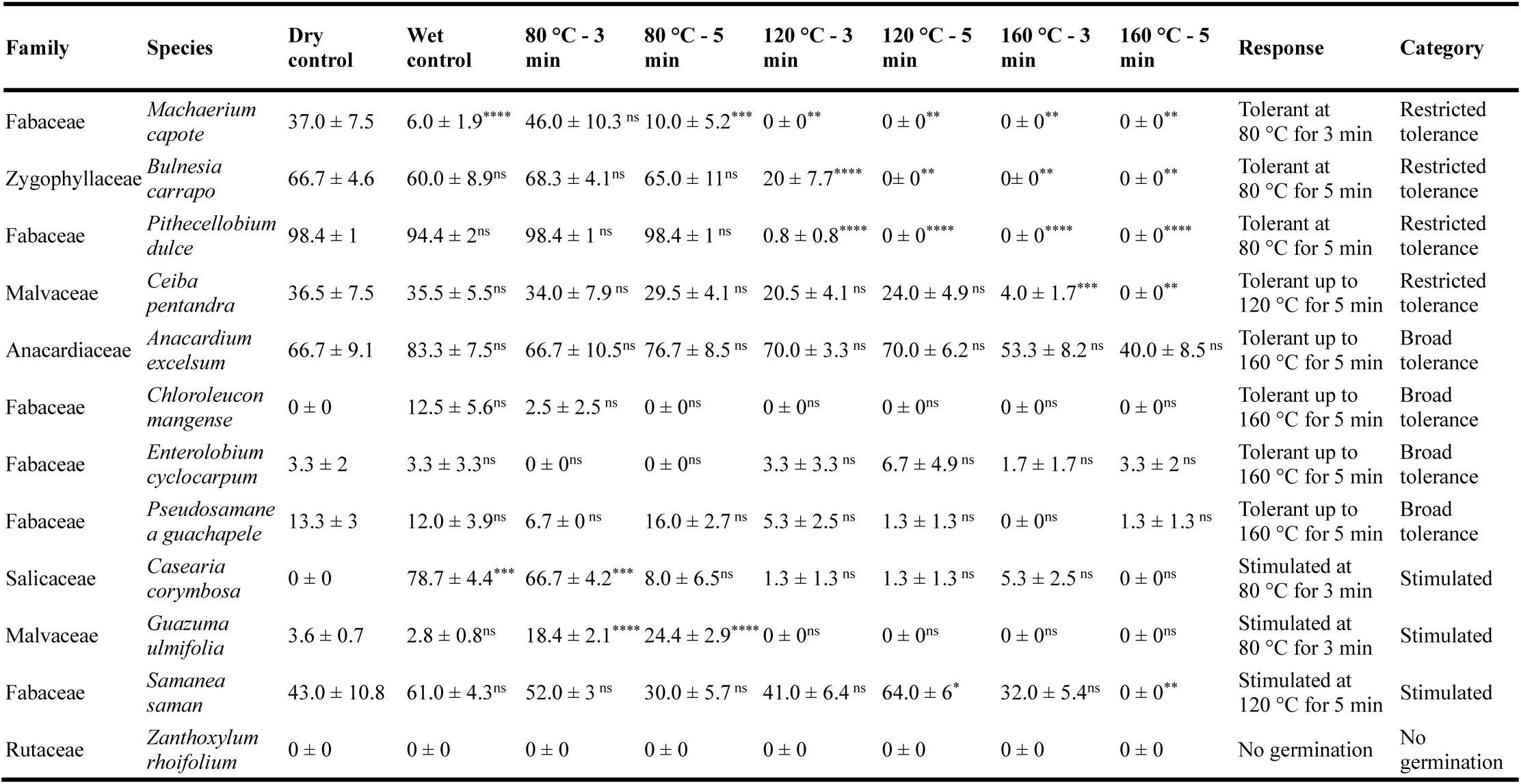
Germination percentage (%, mean ± SE, n = 5 replicates) of the species evaluated under heat shock (HS) treatments. Significance was determined by comparing each treatment with its corresponding dry control. ns = not significant; * p ≤ 0.05; ** p ≤ 0.01; *** p ≤ 0.001; **** p ≤ 0.0001

Among the species with broad tolerance, *Anacardium excelsum*, *Chloroleucon mangense*, *Enterolobium cyclocarpum*, and *Pseudosamanea guachapele* maintained germination responses similar to the control even at 160 °C for both 3 and 5 min (Fig. 1).

Regarding heat-stimulated germination, *Casearia corymbosa* and *Guazuma ulmifolia* showed positive responses at 80 °C, whereas *Samanea saman* showed stimulation at 120 °C for 5 min (Fig. 1). In *C. corymbosa*, germination increased to 66.7% after 3 min at 80 °C relative to the control (OR = 299.04, *p* = 0.0006). In *G. ulmifolia*, germination after 3 and 5 min at 80 °C increased significantly relative to the control (OR = 5.78, *p* < 0.0001; OR = 8.25, *p* < 0.0001, respectively). *Samanea saman* showed a significant increase in germination probability under the 120 °C for 5 min treatment (OR = 2.34, *p* = 0.02).

Post-treatment viability tests complemented the interpretation of the observed germination responses (Online Resource 4). In *Zanthoxylum rhoifolium*, although no germination was recorded under any condition, post-treatment viability decreased under the most intense heat shock treatments. In *M. capote*, *B. carrapo*, *P. dulce*, *A. excelsum*, and *C. corymbosa*, post-treatment viability was zero, indicating that the observed viability was mainly accounted for by seeds that had germinated. In contrast, in *C. pentandra*, *G. ulmifolia*, and *S. saman*, the proportion of germinated and viable seeds was consistent with the germination pattern, with higher values in treatments where tolerance or stimulation was observed.

Although germination in *Chloroleucon mangense*, *Enterolobium cyclocarpum*, and *Pseudosamanea guachapele* was low in several treatments (Table 2), the proportion of germinated and viable seeds was high in several of the lower-intensity treatments, although this response differed among species (Online Resource 4). In *C. mangense*, although seeds exposed to 80 °C for 5 min showed a high proportion of germinated and viable seeds (88.3%), this proportion decreased to 44% in the 120 °C for 3 min treatment and was zero in the treatments from 120 °C for 5 min onward. In *E. cyclocarpum*, this proportion remained at 79.8% even at 160 °C for 5 min. In *P. guachapele*, the proportion remained at 74.7% up to 120 °C for 3 min, with a decrease from 120 °C for 5 min onward.

### Effects of heat shock and smoke on seed germination

Of the 11 species that germinated, smoke significantly increased germination probability only in *Ceiba pentandra*, *Guazuma ulmifolia*, and *Samanea saman*. In the remaining species, significant differences were either absent or attributable to moisture or imbibition effects rather than to smoke itself (Fig. 2; Table 3). In *C. pentandra*, smoke was associated with a significant increase in germination probability at 80 °C for 5 min. Under this treatment, germination reached 48.0%, compared with 29.5% in the equivalent heat treatment without smoke (OR = 2.16, *p* = 0.0072) (Online Resource 5). This pattern was also consistent with the proportion of germinated and viable seeds, which increased from 50.2% without smoke to 62.8% with smoke under the 80 °C for 5 min treatment (Online Resource 6).

**Fig. 2.**
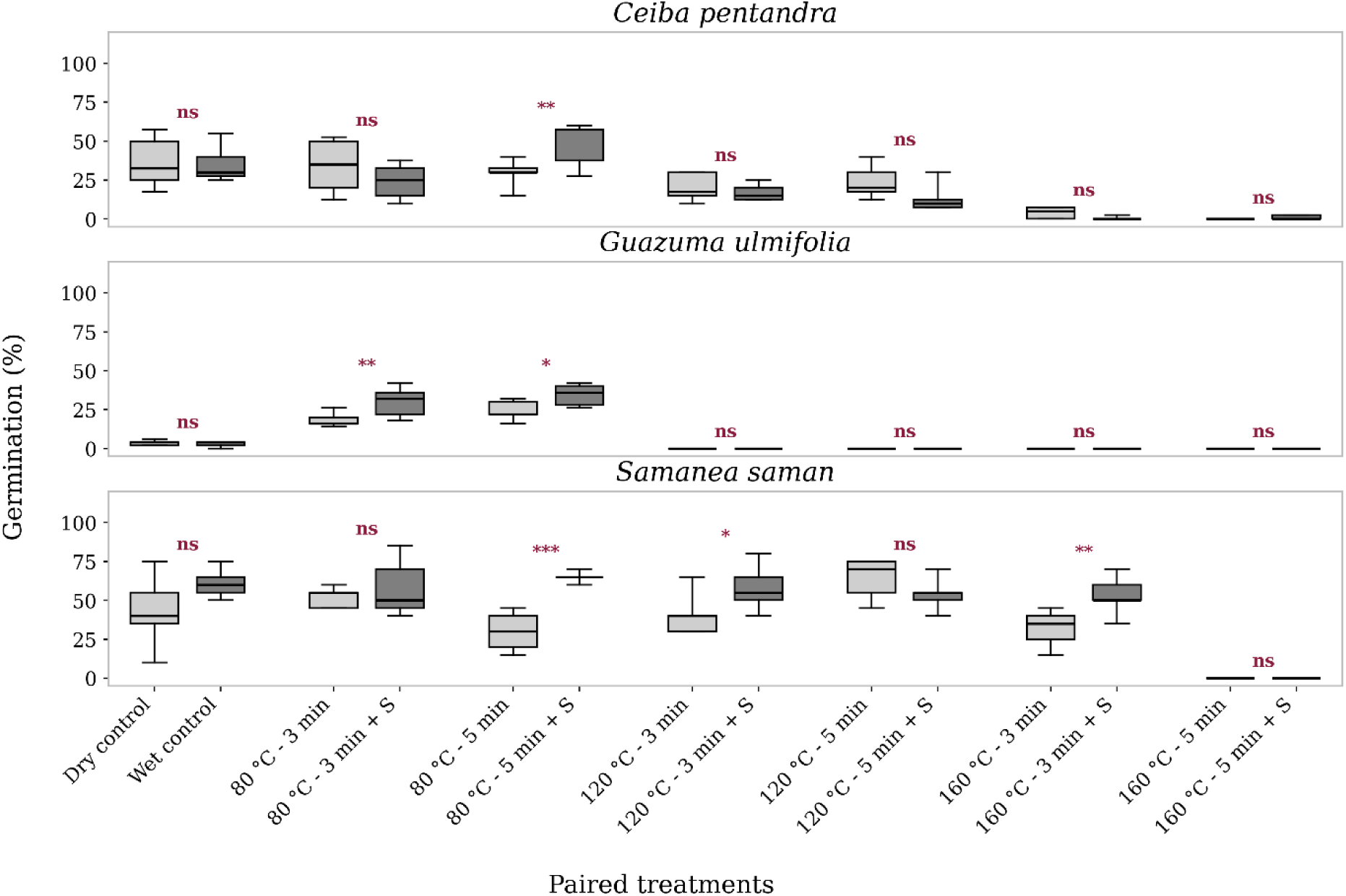
Germination percentage (%) of three species under paired treatments. Paired comparisons are shown between controls (dry vs. wet) and between heat shock treatments (HS) and combined heat shock and smoke treatments (HS+S). Boxes show the interquartile range with the median indicated by the central line; whiskers extend to values within 1.5 times the interquartile range. Significance was determined through pairwise comparisons between the dry and wet controls and between each HS+S treatment and its corresponding heat shock treatment (HS). ns = not significant; * P ≤ 0.05; ** P ≤ 0.01; *** P ≤ 0.001; **** P ≤ 0.0001

**Table 3.** Germination percentage (%, mean ± SE, n = 5) of the species subjected to combined heat shock and smoke treatments (HS+S). Significance was determined through pairwise comparisons between each HS+S treatment and its corresponding heat shock treatment without smoke (HS). ns = not significant; * p ≤ 0.05; ** p ≤ 0.01; *** p ≤ 0.001; **** p ≤ 0.0001. The “Response” column summarizes the overall interpretation of the HS+S effect. For species in which the wet control differed significantly from the dry control, the response was interpreted as being influenced by the effect of moisture

| Family | Species | 80 °C - 3 min + S | 80 °C - 5 min + S | 120 °C - 3 min + S | 120 °C - 5 min + S | 160 °C - 3 min + S | 160 °C - 5 min + S | Response |
| --- | --- | --- | --- | --- | --- | --- | --- | --- |
| Fabaceae | *Machaerium capote* | 10.0 ± 2.7**** | 0 ± 0* | 0 ± 0 | 0 ± 0 | 0 ± 0 | 0 ± 0 | Moisture effect |
| Zygophyllaceae | *Bulnesia carrapo* | 51.7 ± 8.1 | 66.7 ± 9.5 | 25.0 ± 9.5 | 3.3 ± 3.3 | 0 ± 0 | 0 ± 0 | No effect |
| Fabaceae | *Pithecellobium dulce* | 93.6 ± 1 | 93.6 ± 1.6 | 0 ± 0 | 0 ± 0 | 0 ± 0 | 0 ± 0 | No effect |
| Malvaceae | *Ceiba pentandra* | 24.0 ± 5.2 | 48.0 ± 6.5** | 17.0 ± 2.4 | 13.5 ± 4.2 | 0.5 ± 0.5 | 1.0 ± 0.6 | Stimulated by HS+S |
| Anacardiaceae | *Anacardium excelsum* | 60.0 ± 8.5 | 70.0 ± 15.3 | 73.3 ± 6.7 | 63.3 ± 8.2 | 43.3 ± 6.7 | 26.7 ± 8.5 | No effect |
| Fabaceae | *Chloroleucon mangense* | 10.0 ± 4.7 | 2.5 ± 2.5 | 0 ± 0 | 0 ± 0 | 0 ± 0 | 0 ± 0 | No effect |
| Fabaceae | *Enterolobium cyclocarpum* | 3.3 ± 2 | 5.0 ± 3.3 | 5.0 ± 2 | 3.3 ± 2 | 1.7 ± 1.7 | 5.0 ± 3.3 | No effect |
| Fabaceae | *Pseudosamanea guachapele* | 12.0 ± 4.9 | 26.7 ± 6 | 2.7 ± 1.6 | 0 ± 0 | 0 ± 0 | 0 ± 0 | No effect |
| Salicaceae | *Casearia corymbosa* | 72.0 ± 7.4 | 54.7 ± 7.1**** | 5.3 ± 1.3 | 0 ± 0 | 0 ± 0 | 0 ± 0 | Moisture effect |
| Malvaceae | *Guazuma ulmifolia* | 30.0 ± 4.4** | 34.4 ± 3.2 * | 0 ± 0 | 0 ± 0 | 0 ± 0 | 0.0 ± 0 | Stimulated by HS+S |
| Fabaceae | *Samanea saman* | 58.0 ± 8.5 | 65.0 ± 1.6**** | 58.0 ± 6.8* | 54.0 ± 4.8 | 53.0 ± 5.8** | 0 ± 0 | Stimulated by HS+S |
| Rutaceae | *Zanthoxylum rhoifolium* | 0 ± 0 | 0 ± 0 | 0 ± 0 | 0 ± 0 | 0 ± 0 | 0 ± 0 | No effect |

In *G. ulmifolia*, smoke increased germination at 80 °C for 3 and 5 min, reaching 30.0% and 34.4%, respectively, compared with 18.4% and 24.4% in the equivalent heat treatments without smoke (OR = 1.89, *p* = 0.002; OR = 1.62, *p* = 0.01, respectively). In *S. saman*, smoke significantly increased germination probability at 80 °C for 5 min, 120 °C for 3 min, and 160 °C for 3 min (OR = 4.27, *p* < 0.0001; OR = 1.97, *p* = 0.01; OR = 2.37, *p* = 0.003, respectively) (Table 3; Online Resource 5).

In both species, the proportion of germinated and viable seeds was consistent with the germination pattern. In *G. ulmifolia*, combined heat and smoke treatments maintained considerably higher proportions of germinated and viable seeds at 80 °C, reaching 94.6% and 91.1% at 3 and 5 min, respectively, compared with 57.0% and 78.8% in the corresponding heat treatments without smoke. In *S. saman*, smoke was also associated with a higher proportion of germinated and viable seeds in some treatments where germination increased significantly. This proportion increased from 94.4% without smoke to 98.0% with smoke at 120 °C for 3 min, and from 47.9% without smoke to 82.0% with smoke at 160 °C for 3 min. However, this pattern was not consistent across treatments, because at 80 °C for 5 min the values were similar between treatments without and with smoke, at 99.0% and 100.0%, respectively.

### Relationship between functional traits and responses to heat and smoke

Principal component analysis showed that the first two components explained 79.5% of the total variation (PC1 = 51.4% and PC2 = 28.1%) (Fig. 3; trait loadings in Online Resource 7). PC1 was mainly associated with permeability and seed-coat thickness, whereas PC2 was mainly related to shape index.

**Fig. 3.**
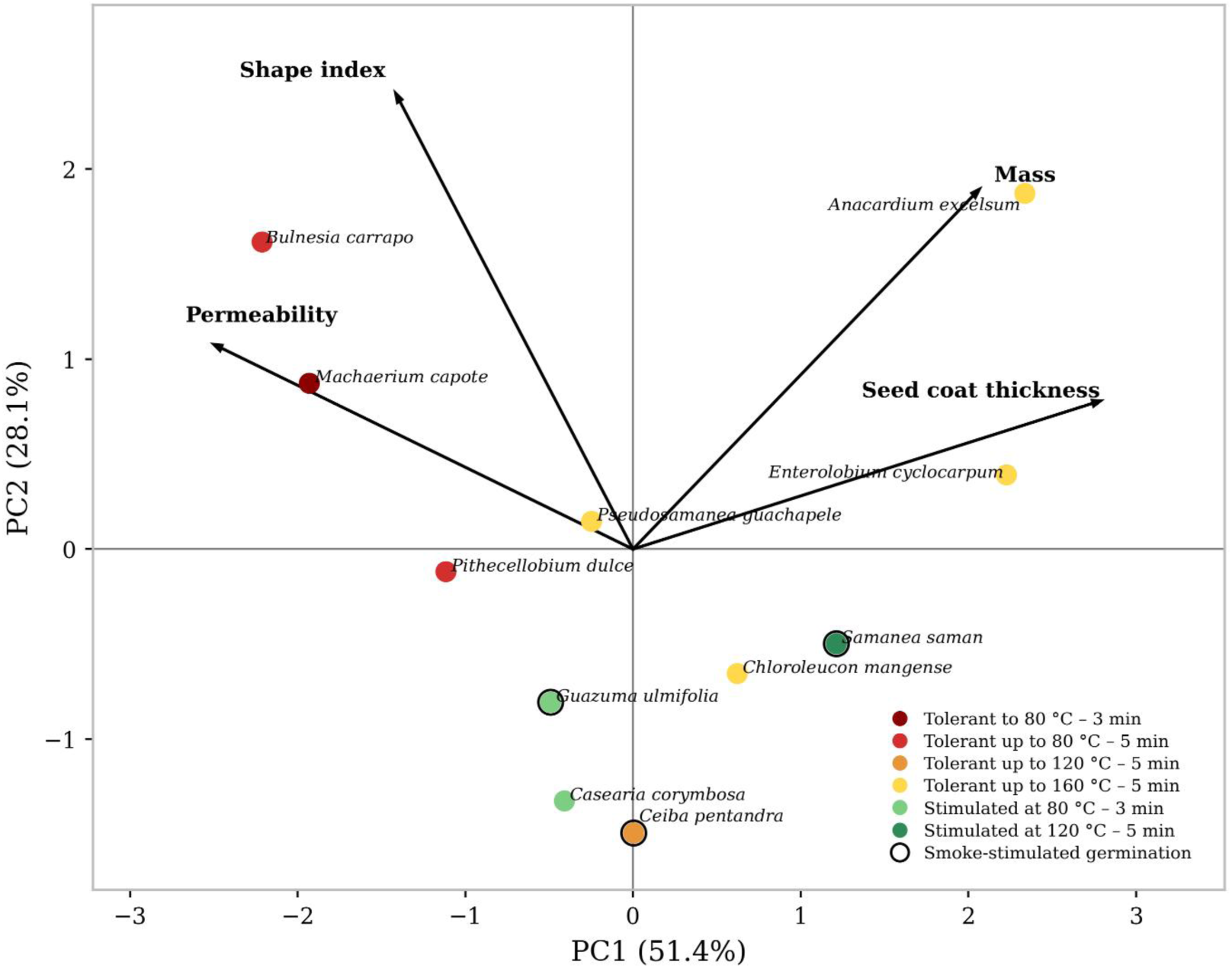
Principal component analysis (PCA) of functional seed traits and their relationship with germination responses to heat shock and combined heat shock and smoke treatments. Arrows indicate trait loadings on the first two principal components. Colors identify the heat-response categories shown in the key, and open circles identify species with smoke-stimulated germination following heat shock

Species showing broad tolerance were positioned toward positive PC1 values, corresponding to seeds with greater mass and thicker seed coats, such as *Anacardium excelsum* and *Enterolobium cyclocarpum*. In contrast, species with restricted tolerance were positioned toward negative PC1 values but positive PC2 values and were characterized by lower seed mass, higher permeability, and higher shape index values, indicating more elongated seeds. This group included *Bulnesia carrapo*, *Machaerium capote*, and *Pithecellobium dulce*. *Ceiba pentandra*, although also classified as having restricted tolerance, occupied a different region of the ordination space, with values close to zero on PC1 and negative values on PC2. Species showing heat-stimulated germination were concentrated in the lower part of the PCA ordination space. This position corresponded to low PC2 values, an axis mainly associated with shape index, indicating less elongated seeds. *Casearia corymbosa* and *Guazuma ulmifolia* were associated with low PC2 values, whereas *Samanea saman* occupied an intermediate position, closer to species with greater seed mass and thicker seed coats.

The three species showing smoke-stimulated germination following heat shock (Fig. 3), *Guazuma ulmifolia, Ceiba pentandra*, and *Samanea saman*, were positioned in the lower region of the PCA, corresponding to negative PC2 values. In addition, *G. ulmifolia* and *C. pentandra* were located toward low or intermediate PC1 values, suggesting a weaker association with greater seed mass and thicker seed coats, whereas S. saman occupied a more positive position along this axis, showing an intermediate association with these traits.

When PC1 was related to the germination response to heat, significant positive relationships were found at 120 °C for 5 min (β = 1.68, R² = 0.52, *p* = 0.012) and at 160 °C for 3 min (β = 1.40, R² = 0.39, *p* = 0.040) (Online Resource 8 and 9). In both treatments, species with positive PC1 values, associated with greater seed mass, thicker seed coats, and lower permeability, showed higher log(OR) values, indicating a smaller reduction in germination response relative to the dry control. At 120 °C for 3 min and 160 °C for 5 min, trends in the same direction were observed, although they were not significant.

For the additional effect of smoke, PC1 showed a significant positive relationship with log(OR) only at 80 °C for 3 min (β = 0.48, R² = 0.41, *p* = 0.034) (Online Resource 10 and 11). Under this treatment, species positioned toward positive PC1 values showed a greater relative effect of smoke on germination compared with their equivalent treatments without smoke. No significant relationships were observed for the remaining temperature and exposure time combinations.

## Discussion

In Colombian TDF, human activities and altered fire regimes may influence regeneration, plant succession, future community composition, and the persistence of species with different functional strategies (Pausas and Keeley 2014; Karna and Jayakumar 2026). In this context, examining seed responses to heat and smoke can help identify species that maintain or increase their germination capacity after disturbance, as well as the functional traits associated with these responses. Previous studies in Colombian TDF have documented contrasting responses to heat shock among species, including tolerance, stimulation, and sensitivity (Cárdenas-Salgado and Pizano 2019; Hernández 2021; Barras et al. 2025). However, whether these responses are also modulated by smoke and by functional seed traits that confer greater or lower protection against fire remains poorly explored. Within this framework, our results show that germination responses among the 11 species that germinated were characterized mainly by tolerance and stimulation in response to heat, and these responses were associated with variation in functional seed traits. In contrast, smoke increased germination following heat shock in only a few species and showed a less consistent functional pattern. *Zanthoxylum rhoifolium*, however, did not germinate under any of the evaluated treatments. Combined dormancy has been reported for this species, involving a thick and impermeable tegmen together with a physiological component and the presence of germination inhibiting compounds (Corrêa et al. 2022). Our fire treatments did not override the dormancy in this case.

Contrary to our first hypothesis, which predicted that the effects of combined heat and smoke treatments would exceed those of heat shock alone, the combination of heat and smoke did not generally produce greater effects than heat alone. This additional effect of smoke increased germination following heat shock in only three of the 11 species studied, *Ceiba pentandra*, *Guazuma ulmifolia*, and *Samanea saman*, which occupied similar positions within the functional trait space. Given that a substantial proportion of seeds did not germinate in the absence of heat and smoke (Fig. 2), heat shock likely acted as the primary mechanism promoting the release of physical dormancy, whereas smoke may have acted as a complementary chemical cue that enhanced processes associated with metabolic activation, respiration, and water uptake, which together could promote germination (Thomas et al. 2003; Light et al. 2009; Keeley et al. 2011; Ramos et al. 2019; Soós et al. 2019; Zirondi et al. 2019; Pausas and Lamont 2022).

Our results partially supported the second hypothesis, in which we expected seeds with low permeability, thick seed coats, greater mass, and a more compact shape to show greater tolerance to heat shock and retain their germination capacity after exposure to high temperatures, whereas more permeable seeds with thinner seed coats, lower mass, and more elongated or flattened shapes were expected to respond positively to smoke and low temperatures (Moreira et al. 2010; Ribeiro et al. 2015; Fichino et al. 2016; Daibes et al. 2019; Hradilová et al. 2019; Monemizadeh et al. 2025). This hypothesis was supported mainly for heat tolerance, as germination responses of TDF seeds were associated with morphological traits related to the physical protection of the embryo. Specifically, the combination of a thicker and less permeable seed coat may have acted as a barrier that protected the embryo and regulated water uptake (Mohamed-Yasseen et al. 1994; Wiggers et al. 2017). Greater seed mass, together with a more compact (rounder) shape, lowers the surface-area-to-volume ratio and may provide a thermal buffering effect, delaying heat transfer to the embryo. Under natural conditions, this morphology may also favor burial and persistence in the soil seed bank, thereby reducing direct exposure to extreme temperatures during fire (Gómez-González et al. 2011; Tavşanoğlu and Çatav 2012; Daibes et al. 2019; Moro et al. 2023). The relationship between this combination of traits and germination was more consistent under the more intense heat treatments.

In contrast, the second part of this hypothesis was not consistently supported because the response to smoke was restricted to only three species. Although these species shared a common functional tendency and occupied similar positions in the trait ordination space, this pattern was not accompanied by the other expected traits, such as higher permeability, lower mass, or thinner seed coats. Instead, our results suggest that under moderate thermal conditions, the additional effect of smoke following heat shock may be greater in species with greater structural protection of the seed. This pattern suggests that responses to smoke may depend more strongly on species-specific characteristics than on a general functional pattern based on the traits evaluated here. Future studies should therefore include a larger number of species representing a broader diversity of functional traits and ecological strategies to strengthen our understanding of the mechanisms regulating germination responses to heat and smoke in this ecosystem.

Although direct records of soil temperatures during wildfires in Colombian TDF are not available, studies from other dry ecosystems provide a useful comparative framework for understanding the thermal conditions that seeds may experience during fire. In seasonally dry tropical forests, most fires are surface fires that primarily consume leaf litter and dead organic material on the forest floor and are generally classified as low to moderate intensity fires (Karna and Jayakumar 2026). Therefore, buried or partially protected seeds may experience lower temperatures than those recorded at the soil surface.

In the Brazilian Cerrado, for example, temperatures at 1 cm depth during fire can approach 80 °C under high fuel loads and exceed 150 °C only under extreme conditions (Daibes et al. 2017; Zupo et al. 2020). Considering that persistent soil seed banks have been documented for several TDF species (Arias Barbosa 2025), our results suggest that seeds buried or partially protected in the soil may survive low to moderate intensity fires.

Indeed, exposure to 80 °C for 3 or 5 min was tolerated by most of the species that germinated and, in some cases, even stimulated germination. At 120 °C, responses began to diverge among species. Some species showed reductions in germination, suggesting an initial thermal threshold for species with restricted tolerance, whereas others maintained or even increased their germination response, as observed in *Samanea saman* at 120 °C for 5 min. This interpretation is consistent with experimental studies from other fire-prone dry ecosystems, where heat shocks between 60 and 100 °C often result in tolerant or stimulated germination responses, whereas temperatures near 150 to 200 °C or prolonged exposure tend to reduce seed viability and germination (Ribeiro et al. 2012; Jaureguiberry and Díaz 2015; Zirondi et al. 2019; Moro et al. 2021; Barras et al. 2025).

Although the predominance of tolerant and heat-stimulated responses among the evaluated species could partly reflect the long history of human influence in Colombian TDF, because many of these species are widely used, managed, or favored by human communities (Arroyo-Rodríguez et al. 2015), our results indicate that variation in germination responses to heat was associated with functional seed traits. In particular, broad heat tolerance was associated with lower permeability, thicker seed coats, and greater seed mass, whereas heat-stimulated germination showed a different pattern, more closely related to variation in seed shape. This suggests that other species, regardless of their degree of association with human use, may show comparable responses to heat exposure if they share these functional traits. *Anacardium excelsum* illustrates this pattern because, despite being associated with intermediate successional forests and riparian environments where fire is less frequent (Yepes-Quintero et al. 2007), it showed broad heat tolerance associated with its large seed mass and intermediate permeability, traits that may contribute substantially to seed survival under heat exposure.

These results have several implications for the management of natural regeneration and post-fire restoration in TDF landscapes subject to recurrent fires. Species whose seeds show broad germination tolerance to heat, associated with lower permeability, thicker seed coats, and greater mass, as well as species with heat-stimulated germination, may have a greater capacity to maintain germination potential following fire-related heat exposure. These characteristics may be particularly relevant in transition zones between productive systems and forests, where fires occur more frequently because of agricultural burning, pasture management, or accidental fires associated with litter accumulation and removal (Ramírez 2020). In contrast, species with restricted tolerance or sensitivity to certain levels of heat, associated with more permeable, elongated, or flattened seeds and thinner seed coats, may be more vulnerable to reductions in germination following intense heat exposure. Such species may therefore require greater attention in enrichment strategies within forest interiors, reserve areas, and conservation zones to maintain functional diversity and prevent regeneration from becoming dominated by a single set of functional traits.

Complementarily, the observed heat tolerance and stimulation suggest that prescribed burning could be explored, within future integrated fire management frameworks, as a potential tool for reducing fuel loads in transition zones between productive systems and forests and for lowering the risk of more severe fires that could affect the soil seed bank. However, the implementation of such practices would require careful consideration of the thermal tolerance thresholds of the most sensitive species to avoid negative effects on natural regeneration, seed bank persistence, and the functional composition of TDF.

## Acknowledgements

We thank Universidad Distrital Francisco José de Caldas, particularly the Wood Laboratory, for its support during the experimental phase of this study. We also acknowledge the University of Sussex for its institutional support and collaboration during the development of this research. Finally, we thank Juan Garibello for his support with the statistical analyses.

## Author contributions

N.A.D.B. contributed to conceptualization, methodology, investigation, data curation, formal analysis, visualization, and writing of the original draft. M.C.C. contributed to conceptualization, methodology, supervision, formal analysis, and writing—review and editing. A.P.R. contributed to conceptualization, methodology, supervision, project administration, interpretation of results, and writing—review and editing.

## Funding

This research was funded by the Sussex International Development Challenge Fund through the project “Fire-smart restoration for resilient livelihoods in tropical dry forest landscapes,” developed in collaboration between Universidad Distrital Francisco José de Caldas and the University of Sussex.

## Conflicts of interest

The authors declare that they have no conflict of interest.

## Availability of data and material

The datasets generated and/or analyzed during the current study are available from the corresponding author on reasonable request.

## Code availability

The R code used for the statistical analyses is available from the corresponding author on reasonable request.

## Notes

### Competing Interest Statement

The authors have declared no competing interest.

