## Supplementary Material for "Effects of Heat Shock and Smoke on the Germination of Woody Species from Tropical Dry Forests"

**Online Resource 1.** Species, fruit type, seed storage behavior, collection date and season, experiment date, and number of seeds per replicate used in the germination treatments

| Family | Species | Fruit type | Seed storage behavior | Collection date | Collection season | Experiment date | Seeds per replicate |
|---|---|---|---|---|---|---|---|
| Fabaceae | *Machaerium capote* | Samaroid legume (Meléndez González 2009) | Orthodox (Meléndez González 2009) | 22-Mar-2025 | First rainy season or transition to the rainy season | 27-Aug-2025 | 20 |
| Zygophyllaceae | *Bulnesia carrapo* | Winged schizocarp (Gómez Gamarra 2017) | Orthodox (Gómez Gamarra 2017) | 13-Apr-2025 | First rainy season | 03-May-2025 | 12 |
| Fabaceae | *Pithecellobium dulce* | Dehiscent legume (Vargas-Figueroa 2015) | Orthodox (Vargas-Figueroa 2015) | 23-Mar-2025 | First rainy season or transition to the rainy season | 08-Apr-2025 | 25 |
| Malvaceae | *Ceiba pentandra* | Dehiscent capsule (Chan et al. 2023) | Orthodox (Chan et al. 2023) | 15-Apr-2025 | First rainy season | 08-May-2025 | 40 |
| Anacardiaceae | *Anacardium excelsum* | Reniform drupe (Sautu et al. 2006) | Orthodox (Sautu et al. 2006) | 24-Mar-2025 | First rainy season or transition to the rainy season | 29-Mar-2025 | 6 |
| Fabaceae | *Chloroleucon mangense* | Dehiscent legume (Rodríguez-Trejo and Guizar-Nolazco 2016) | Orthodox (Rodríguez-Trejo and Guizar-Nolazco 2016) | 03-Nov-2025 | Second rainy season | 06-Nov-2025 | 8 |
| Fabaceae | *Enterolobium cyclocarpum* | Indehiscent legume (Arceo-Gómez et al. 2022) | Orthodox (Arceo-Gómez et al. 2022) | 22-Mar-2025 | First rainy season or transition to the rainy season | 22-Aug-2025 | 12 |
| Fabaceae | *Pseudosamanea guachapele* | Dehiscent legume (Rico Arce et al. 2008) | Orthodox (Rico Arce et al. 2008) | 13-Apr-2025 | First rainy season | 11-Oct-2025 | 15 |
| Salicaceae | *Casearia corymbosa* | Dehiscent capsule (Pool and Smith 2001) | Recalcitrant (Farnsworth 2000) | 14-Oct-2025 | Second rainy season | 22-Oct-2025 | 15 |
| Malvaceae | *Guazuma ulmifolia* | Woody indehiscent capsule (Paiva Sobrinho and Siqueira 2008) | Orthodox (Muñoz et al. 2004) | 07-Aug-2025 | Dry season | 27-Aug-2025 | 50 |
| Fabaceae | *Samanea saman* | Indehiscent legume (Elevitch and Wilkinson 2006) | Orthodox (Vargas-Figueroa 2015) | 13-Apr-2025 | First rainy season | 11-Oct-2025 | 20 |
| Rutaceae | *Zanthoxylum rhoifolium* | Dry dehiscent follicle (Silva and Paoli 2000) | Orthodox (Corrêa et al. 2021) | 14-Apr-2025 | First rainy season | 22-Aug-2025 | 50 |

**Online Resource 2.** Functional seed traits (mean ± SE, n = 50), including permeability, seed coat thickness, seed mass, and shape index

| Family | Species | Permeability (% ± SE) | Permeability class | Seed coat thickness (µm ± SE) | Seed mass (g ± SE) | Length (mm ± SE) | Width (mm ± SE) | Thickness (mm ± SE) | Shape index (mean ± SE) |
|---|---|---|---|---|---|---|---|---|---|
| Fabaceae | *Machaerium capote* | 113.73 ± 6.89 | Permeable | 40 ± 2 | 0.04 ± 0.00 | 9.52 ± 0.13 | 3.75 ± 0.06 | 1.53 ± 0.03 | 0.19 ± 0.001 |
| Zygophyllaceae | *Bulnesia carrapo* | 160.43 ± 9.38 | Permeable | 168 ± 7 | 0.16 ± 0.01 | 17.14 ± 0.25 | 11.13 ± 0.17 | 1.81 ± 0.04 | 0.20 ± 0.001 |
| Fabaceae | *Pithecellobium dulce* | 119.37 ± 4.82 | Permeable | 190 ± 5 | 0.08 ± 0.00 | 7.76 ± 0.09 | 6.32 ± 0.07 | 3.10 ± 0.06 | 0.10 ± 0.003 |
| Malvaceae | *Ceiba pentandra* | 58.71 ± 5.47 | Permeable | 196 ± 7 | 0.04 ± 0.00 | 5.26 ± 0.04 | 4.27 ± 0.05 | 3.68 ± 0.04 | 0.03 ± 0.001 |
| Anacardiaceae | *Anacardium excelsum* | 46.97 ± 5.25 | Permeable | 677 ± 18 | 2.47 ± 0.06 | 31.17 ± 0.22 | 18.36 ± 0.12 | 10.91 ± 0.13 | 0.11 ± 0.001 |
| Fabaceae | *Chloroleucon mangense* | 30.68 ± 7.05 | Permeable | 481 ± 8 | 0.04 ± 0.00 | 5.80 ± 0.06 | 3.83 ± 0.05 | 2.45 ± 0.06 | 0.09 ± 0.003 |
| Fabaceae | *Enterolobium cyclocarpum* | 18.34 ± 4.12 | Impermeable | 938 ± 15 | 0.79 ± 0.02 | 17.44 ± 0.11 | 10.54 ± 0.11 | 6.76 ± 0.07 | 0.10 ± 0.001 |
| Fabaceae | *Pseudosamanea guachapele* | 10.38 ± 1.96 | Impermeable | 208 ± 7 | 0.04 ± 0.00 | 9.12 ± 0.06 | 4.37 ± 0.09 | 1.60 ± 0.02 | 0.18 ± 0.001 |
| Salicaceae | *Casearia corymbosa* | 60.23 ± 5.88 | Permeable | 58 ± 2 | 0.07 ± 0.00 | 6.70 ± 0.11 | 5.00 ± 0.06 | 4.07 ± 0.11 | 0.05 ± 0.003 |
| Malvaceae | *Guazuma ulmifolia* | 74.37 ± 8.39 | Permeable | 189 ± 6 | 0.004 ± 0.000 | 2.76 ± 0.02 | 1.75 ± 0.03 | 1.34 ± 0.03 | 0.07 ± 0.002 |
| Fabaceae | *Samanea saman* | 14.95 ± 3.23 | Impermeable | 613 ± 11 | 0.21 ± 0.01 | 11.52 ± 0.11 | 6.90 ± 0.06 | 4.87 ± 0.06 | 0.09 ± 0.002 |
| Rutaceae | *Zanthoxylum rhoifolium* | 28.89 ± 2.00 | Permeable | 749 ± 20 | 0.02 ± 0.00 | 3.74 ± 0.03 | 3.34 ± 0.03 | 2.75 ± 0.04 | 0.02 ± 0.001 |

**Online Resource 3.** Results of the generalized linear models: odds ratios and *p* values for comparisons of the heat shock treatments and the wet control with the dry control. Values are odds ratios, with *p* values in parentheses

| Family | Species | Wet control | 80 °C - 3 min | 80 °C - 5 min | 120 °C - 3 min | 120 °C - 5 min | 160 °C - 3 min | 160 °C - 5 min |
|---|---|---|---|---|---|---|---|---|
| Fabaceae | *Machaerium capote* | 0.116 (<0.0001) | 1.445 (0.6423) | 0.196 (0.0002) | 0.008 (0.0059) | 0.008 (0.0059) | 0.008 (0.0059) | 0.008 (0.0059) |
| Zygophyllaceae | *Bulnesia carrapo* | 0.754 (0.9150) | 1.077 (0.9988) | 0.930 (0.9989) | 0.130 (<0.0001) | 0.004 (0.0012) | 0.004 (0.0012) | 0.004 (0.0012) |
| Fabaceae | *Pithecellobium dulce* | 0.3200 (0.4769) | 1.0000 (1.0000) | 1.0000 (1.0000) | 0.0002 (<0.0001) | 0.0001 (<0.0001) | 0.0001 (<0.0001) | 0.0001 (<0.0001) |
| Malvaceae | *Ceiba pentandra* | 0.9575 (0.9997) | 0.8962 (0.9932) | 0.7280 (0.8386) | 0.4486 (0.1444) | 0.5494 (0.3744) | 0.0811 (0.0002) | 0.0215 (0.0015) |
| Anacardiaceae | *Anacardium excelsum* | 2.3750 (0.5567) | 1.0000 (1.0000) | 1.6050 (0.8862) | 1.1590 (0.9965) | 1.1590 (0.9965) | 0.5830 (0.7971) | 0.3460 (0.2281) |
| Fabaceae | *Chloroleucon mangense* | 12.5500 (0.3897) | 3.0800 (0.9366) | 1.0000 (1.0000) | 1.0000 (1.0000) | 1.0000 (1.0000) | 1.0000 (1.0000) | 1.0000 (1.0000) |
| Fabaceae | *Enterolobium cyclocarpum* | 1.0000 (1.0000) | 0.1930 (0.7814) | 0.1930 (0.7814) | 1.0000 (1.0000) | 1.8640 (0.9089) | 0.5900 (0.9746) | 1.0000 (1.0000) |
| Fabaceae | *Pseudosamanea guachapele* | 0.8912 (0.9976) | 0.4867 (0.6322) | 1.2280 (0.9817) | 0.3926 (0.4475) | 0.1256 (0.1101) | 0.0413 (0.1533) | 0.1256 (0.1101) |
| Salicaceae | *Casearia corymbosa* | 544.5200 (0.0001) | 299.0400 (0.0006) | 14.1200 (0.3269) | 3.0400 (0.9367) | 3.0400 (0.9367) | 9.5000 (0.5025) | 1.0000 (1.0000) |
| Malvaceae | *Guazuma ulmifolia* | 0.7830 (0.9757) | 5.7803 (<0.0001) | 8.2501 (<0.0001) | 0.0507 (0.1997) | 0.0507 (0.1997) | 0.0507 (0.1997) | 0.0507 (0.1997) |
| Fabaceae | *Samanea saman* | 2.0581 (0.0687) | 1.4309 (0.6528) | 0.5719 (0.2742) | 0.9220 (0.9957) | 2.3359 (0.0212) | 0.6272 (0.4425) | 0.0066 (0.0032) |

**Online Resource 4.** Germination plus viable seed percentage (%, mean ± SE, n = 5) of species subjected to heat shock treatments. Species for which post-treatment viability of ungerminated seeds was zero were not included in the table

| Family | Species | Dry control | Wet control | 80 °C - 3 min | 80 °C - 5 min | 120 °C - 3 min | 120 °C - 5 min | 160 °C - 3 min | 160 °C - 5 min |
|---|---|---|---|---|---|---|---|---|---|
| Malvaceae | *Ceiba pentandra* | 36.5 ± 7.5 | 62.2 ± 10.2 | 46.0 ± 3.0 | 50.2 ± 7.3 | 35.2 ± 8.1 | 28.3 ± 3.6 | 7.1 ± 2.2 | 12.5 ± 6.8 |
| Fabaceae | *Chloroleucon mangense* | 84.2 ± 9.2 | 90.0 ± 4.7 | 89.2 ± 4.9 | 88.3 ± 3.1 | 44.0 ± 11.2 | 0.0 ± 0.0 | 0.0 ± 0.0 | 0.0 ± 0.0 |
| Fabaceae | *Enterolobium cyclocarpum* | 67.7 ± 8.1 | 80.8 ± 4.9 | 67.1 ± 4.3 | 58.6 ± 4.7 | 82.8 ± 5.9 | 76.0 ± 5.2 | 69.1 ± 5.4 | 79.8 ± 3.3 |
| Fabaceae | *Pseudosamanea guachapele* | 100.0 ± 0.0 | 97.9 ± 2.1 | 96.0 ± 4.0 | 100.0 ± 0.0 | 74.7 ± 6.5 | 20.7 ± 3.3 | 0.0 ± 0.0 | 1.3 ± 1.3 |
| Malvaceae | *Guazuma ulmifolia* | 74.7 ± 3.4 | 81.6 ± 3.3 | 57.0 ± 3.6 | 78.8 ± 2.6 | 20.7 ± 15.1 | 0.0 ± 0.0 | 0.0 ± 0.0 | 0.0 ± 0.0 |
| Fabaceae | *Samanea saman* | 100.0 ± 0.0 | 97.0 ± 2.0 | 99.0 ± 1.0 | 99.0 ± 1.0 | 94.4 ± 2.3 | 73.0 ± 7.5 | 47.9 ± 7.8 | 0.0 ± 0.0 |
| Rutaceae | *Zanthoxylum rhoifolium* | 70.0 ± 5.7 | 62.9 ± 4.2 | 47.1 ± 14.0 | 20.0 ± 5.7 | 0.0 ± 0.0 | 0.0 ± 0.0 | 0.0 ± 0.0 | 0.0 ± 0.0 |

**Online Resource 5.** Results of the generalized linear models: odds ratios and p values for comparisons of the combined heat shock and smoke treatments with the corresponding heat shock treatments without smoke. Values are odds ratios, with *p* values in parentheses

| **Family** | **Species** | **80 °C - 3 min + S** | **80 °C - 5 min + S** | **120 °C - 3 min + S** | **120 °C - 5 min + S** | **160 °C - 3 min + S** | **160 °C - 5 min + S** |
|---|---|---|---|---|---|---|---|
| Fabaceae | *Machaerium capote* | 0.1360 | 0.0429 | 1.0000 | 1.0000 | 1.0000 | 1.0000 |
| | | (<0.0001) | (0.0312) | (1.0000) | (1.0000) | (1.0000) | (1.0000) |
| Zygophyllaceae | *Bulnesia carrapo* | 0.5020 | 1.0750 | 1.3220 | 5.1710 | 1.0000 | 1.0000 |
| | | (0.0685) | (0.8501) | (0.5224) | (0.2962) | (1.0000) | (1.0000) |
| Fabaceae | *Pithecellobium dulce* | 0.2800 | 0.2800 | 0.3310 | 1.0000 | 1.0000 | 1.0000 |
| | | (0.0827) | (0.0827) | (0.5010) | (1.0000) | (1.0000) | (1.0000) |
| Malvaceae | *Ceiba pentandra* | 0.6240 | 2.1580 | 0.8040 | 0.5140 | 0.3220 | 1.8180 |
| | | (0.1188) | (0.0072) | (0.5288) | (0.0611) | (0.1900) | (0.5869) |
| Anacardiaceae | *Anacardium excelsum* | 0.7580 | 0.7220 | 1.1700 | 0.7490 | 0.6780 | 0.5590 |
| | | (0.6053) | (0.5761) | (0.7832) | (0.5979) | (0.4540) | (0.2933) |
| Fabaceae | *Chloroleucon mangense* | 3.2500 | 3.0800 | 1.0000 | 1.0000 | 1.0000 | 1.0000 |
| | | (0.2307) | (0.5007) | (1.0000) | (1.0000) | (1.0000) | (1.0000) |
| Fabaceae | *Enterolobium cyclocarpum* | 5.1710 | 7.3650 | 1.4240 | 0.5370 | 1.0000 | 1.4240 |
| | | (0.2962) | (0.1935) | (0.6793) | (0.4462) | (1.0000) | (0.6793) |
| Fabaceae | *Pseudosamanea guachapele* | 1.8310 | 1.8760 | 0.5400 | 0.3290 | 1.0000 | 0.3290 |
| | | (0.2853) | (0.1210) | (0.4482) | (0.5009) | (1.0000) | (0.5009) |
| Salicaceae | *Casearia corymbosa* | 1.2800 | 12.8620 | 3.1260 | 0.3290 | 0.1050 | 1.0000 |
| | | (0.4860) | (<0.0001) | (0.2369) | (0.5009) | (0.1359) | (1.0000) |
| Malvaceae | *Guazuma ulmifolia* | 1.8900 | 1.6200 | 1.0000 | 1.0000 | 1.0000 | 1.0000 |
| | | (0.0028) | (0.0150) | (1.0000) | (1.0000) | (1.0000) | (1.0000) |
| Fabaceae | *Samanea saman* | 1.2720 | 4.2650 | 1.9730 | 0.6630 | 2.3740 | 1.0000 |
| | | (0.3988) | (<0.0001) | (0.0178) | (0.1555) | (0.0032) | (1.0000) |

**Online Resource 6.** Germination plus viable seed percentage (%, mean ± SE, n = 5) of species subjected to combined heat shock and smoke treatments (HS+S). Species for which post-treatment viability of ungerminated seeds was zero were not included in the table

| Family | Species | 80 °C - 3 min + S | 80 °C - 5 min + S | 120 °C - 3 min + S | 120 °C - 5 min + S | 160 °C - 3 min + S | 160 °C - 5 min + S |
|---|---|---|---|---|---|---|---|
| Malvaceae | *Ceiba pentandra* | 58.9 ± 5.0 | 62.8 ± 2.9 | 32.2 ± 10.2 | 34.1 ± 8.2 | 0.5 ± 0.5 | 1.0 ± 0.6 |
| Fabaceae | *Chloroleucon mangense* | 91.5 ± 3.6 | 80.5 ± 4.0 | 20.5 ± 3.8 | 9.0 ± 6.0 | 0.0 ± 0.0 | 0.0 ± 0.0 |
| Fabaceae | *Enterolobium cyclocarpum* | 81.7 ± 8.0 | 66.1 ± 5.9 | 72.4 ± 9.2 | 75.1 ± 6.6 | 63.3 ± 5.9 | 61.3 ± 7.6 |
| Fabaceae | *Pseudosamanea guachapele* | 100.0 ± 0.0 | 93.4 ± 4.1 | 42.8 ± 5.6 | 32.6 ± 7.1 | 9.1 ± 4.1 | 0.0 ± 0.0 |
| Malvaceae | *Guazuma ulmifolia* | 94.6 ± 1.8 | 91.1 ± 2.5 | 0.0 ± 0.0 | 0.0 ± 0.0 | 2.5 ± 2.5 | 0.0 ± 0.0 |
| Fabaceae | *Samanea saman* | 99.0 ± 1.0 | 100.0 ± 0.0 | 98.0 ± 1.2 | 82.0 ± 3.0 | 82.0 ± 3.4 | 1.0 ± 1.0 |
| Rutaceae | *Zanthoxylum rhoifolium* | 40.0 ± 11.6 | 27.1 ± 8.3 | 0.0 ± 0.0 | 0.0 ± 0.0 | 0.0 ± 0.0 | 0.0 ± 0.0 |

**Online Resource 7.** Loadings of functional seed traits on the first two principal components (PC1 and PC2) of the principal component analysis (PCA)

| Functional seed trait | PC1 | PC2 |
|---|---|---|
| Seed-coat thickness | 0.888 | 0.249 |
| Permeability | -0.795 | 0.343 |
| Seed mass | 0.657 | 0.603 |
| Shape index | -0.450 | 0.764 |
| **Explained variance (%)** | **51.4** | **28.1** |

**Online Resource 8.** Relationship between the functional seed trait gradient (PC1) and germination response to heat shock. **Note:** β represents the slope of the linear model log(OR) ~ PC1; SE = standard error; $R^2$ = coefficient of determination; ρ = Spearman's correlation coefficient. Associations were considered significant when $p < 0.05$

| **Treatment** | **β PC1** | **SE** | **t** | **$R^2$** | **Adjusted $R^2$** | **Regression *p*** | **Spearman's ρ** | **Spearman's *p*** | **Interpretation** |
|---|---|---|---|---|---|---|---|---|---|
| 80 °C – 3 min | -0.298 | 0.430 | -0.692 | 0.051 | -0.055 | 0.506 | -0.309 | 0.355 | Not significant |
| 80 °C – 5 min | -0.098 | 0.305 | -0.320 | 0.011 | -0.099 | 0.756 | -0.100 | 0.770 | Not significant |
| 120 °C – 3 min | 1.140 | 0.518 | 2.190 | 0.348 | 0.276 | 0.056 | 0.718 | 0.013 | Positive trend; significant according to Spearman's correlation |
| 120 °C – 5 min | 1.680 | 0.538 | 3.120 | 0.519 | 0.466 | 0.012 | 0.745 | 0.008 | Significant positive association |
| 160 °C – 3 min | 1.400 | 0.584 | 2.390 | 0.389 | 0.321 | 0.040 | 0.673 | 0.023 | Significant positive association |
| 160 °C – 5 min | 1.170 | 0.563 | 2.080 | 0.324 | 0.249 | 0.067 | 0.527 | 0.096 | Positive trend |

**Online Resource 9.** Relationship between the first principal component (PC1) of functional seed traits and germination response to heat shock, expressed as log(OR) relative to the dry control, for the 120 °C for 5 min and 160 °C for 3 min treatments. The blue line represents the fitted linear model, and the gray band represents the 95% confidence interval

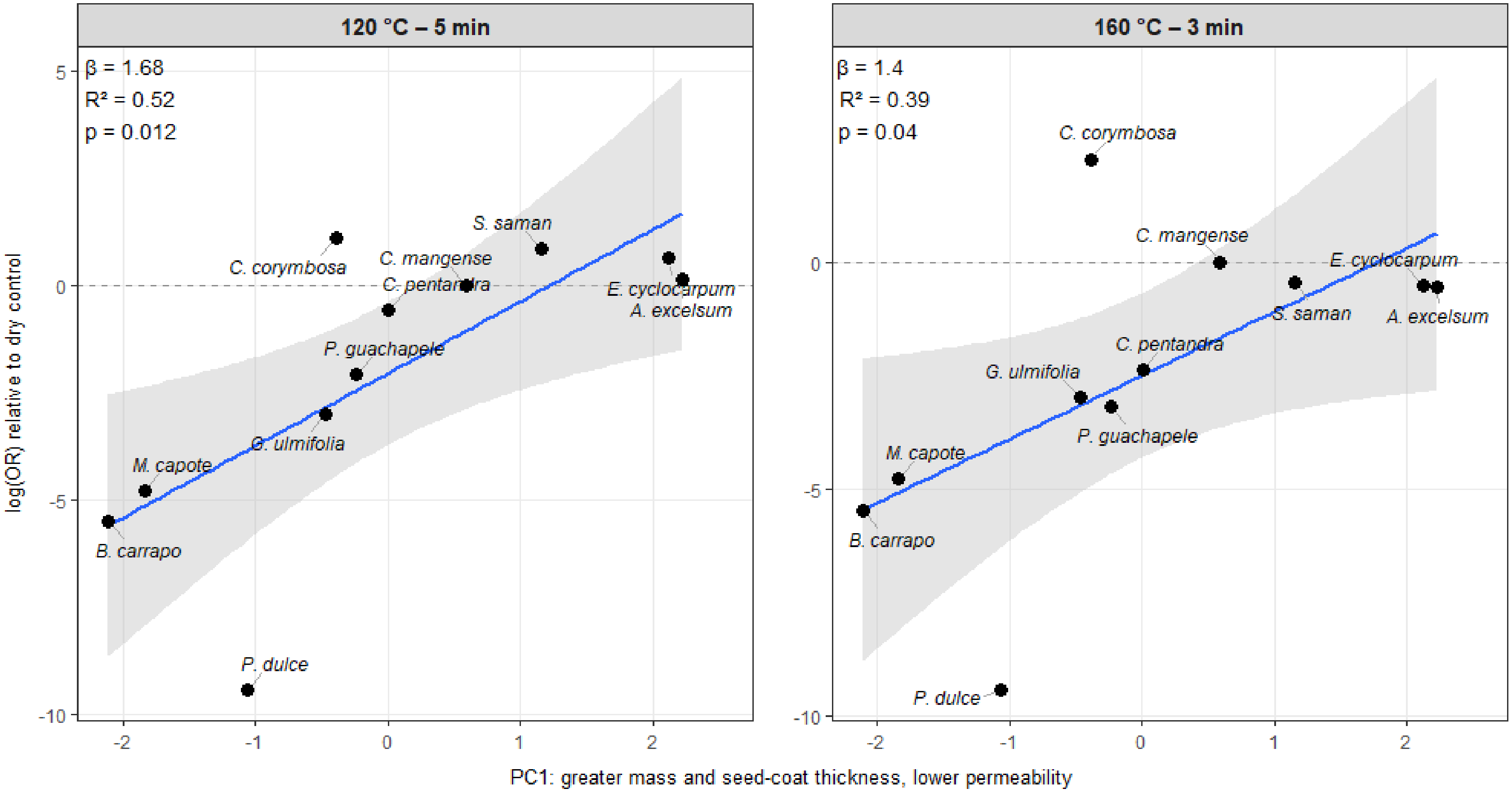

**Online Resource 10.** Relationship between the functional seed trait gradient (PC1) and the effect of smoke on germination following heat shock treatments. Note: β represents the slope of the linear model log(OR) ~ PC1; SE = standard error; $R^2$ = coefficient of determination; ρ = Spearman's correlation coefficient. Associations were considered significant when $p < 0.05$

| Treatment | β PC1 | SE | t | $R^2$ | Adjusted $R^2$ | Regression *p* | Spearman's ρ | Spearman's *p* | Interpretation |
|---|---|---|---|---|---|---|---|---|---|
| 80 °C – 3 min | 0.476 | 0.190 | 2.500 | 0.411 | 0.345 | 0.0336 | 0.555 | 0.0767 | Significant positive association in the regression; Spearman's correlation not significant |
| 80 °C – 5 min | 0.559 | 0.317 | 1.760 | 0.256 | 0.174 | 0.1120 | 0.491 | 0.1250 | Not significant |
| 120 °C – 3 min | 0.106 | 0.135 | 0.784 | 0.064 | -0.040 | 0.4530 | 0.327 | 0.3260 | Not significant |
| 120 °C – 5 min | -0.254 | 0.153 | -1.660 | 0.234 | 0.149 | 0.1320 | -0.418 | 0.2010 | Not significant |
| 160 °C – 3 min | 0.049 | 0.187 | 0.263 | 0.008 | -0.103 | 0.7990 | -0.082 | 0.8110 | Not significant |
| 160 °C – 5 min | -0.014 | 0.104 | -0.131 | 0.002 | -0.109 | 0.8990 | -0.128 | 0.7070 | Not significant |

**Online Resource 11.** Relationship between the first principal component (PC1) of functional seed traits and the additive effect of smoke on germination, expressed as log(OR) for the comparison of treatments with smoke versus without smoke, for the 80 °C for 3 min treatment. The blue line represents the fitted linear model, and the gray band represents the 95% confidence interval

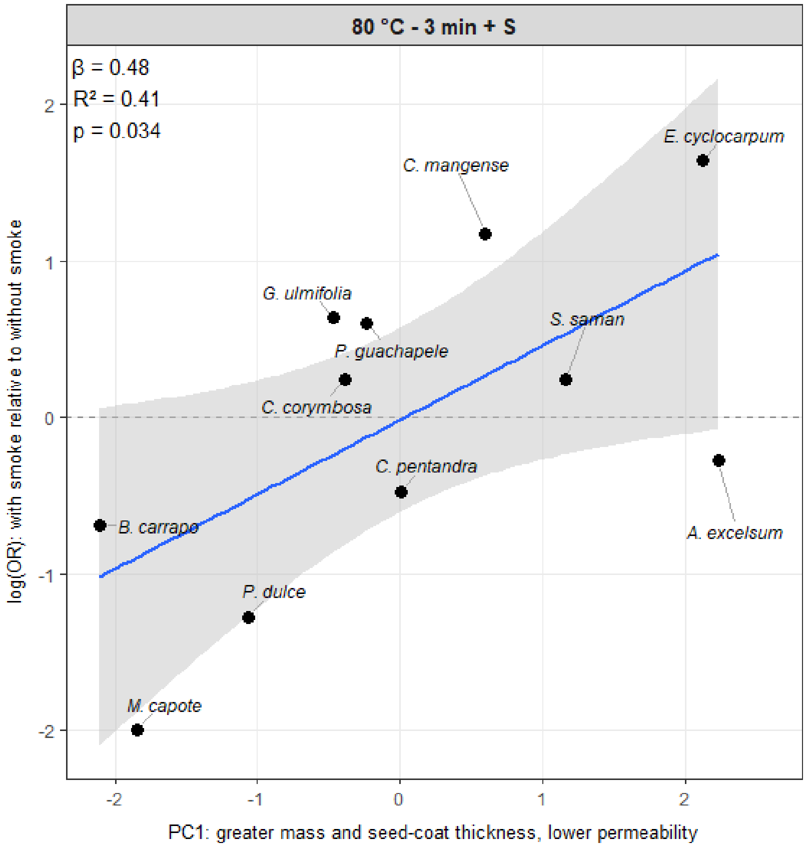
